# The sensory code within sense of time

**DOI:** 10.1101/2021.07.29.454157

**Authors:** S. Reinartz, A. Fassihi, L. Paz, F. Pulecchi, M. Gigante, M.E. Diamond

**Author notes:** These authors contributed equally to this work (S. Reinartz, A. Fassihi).

## Abstract

Sensory experiences are accompanied by the perception of the passage of time; a cell phone vibration, for instance, is sensed as brief or long. The neuronal mechanisms underlying the perception of elapsed time remain unknown^1^. Recent work agrees on a role for cortical processing networks^2,3^, however the causal function of sensory cortex in time perception has not yet been specified. We hypothesize that the mechanisms for time perception are embedded within primary sensory cortex and are thus governed by the basic rules of sensory coding. By recording and optogenetically modulating neuronal activity in rat vibrissal somatosensory cortex, we find that the percept of stimulus duration is dilated and compressed by optogenetic excitation and inhibition, respectively, during stimulus delivery. A second set of rats judged the intensity of tactile stimuli; here, optogenetic excitation amplified the intensity percept, demonstrating sensory cortex to be the common gateway to both time and stimulus feature processing. The coding algorithms for sensory features are well established^4–10^. Guided by these algorithms, we formulated a 3-stage model beginning with the membrane currents evoked by vibrissal and optogenetic drive and culminating in the representation of perceived time; this model successfully replicated rats’ choices. Our finding that stimulus coding is intrinsic to sense of time disagrees with dedicated pacemaker-accumulator operation models^11–13^, where sensory input acts only to trigger the onset and offset of the timekeeping process. Time perception is thus as deeply intermeshed within the sensory processing pathway as is the sense of touch itself^14,15^ and can now be treated through the computational language of sensory coding. The model presented here readily generalizes to humans^14,16^ and opens up new approaches to understanding the time misperception at the core of numerous neurological conditions^17,18^.

---

The subjective experience of an external stimulus has a dual nature – the feeling of the physical features of the sensory input and, in parallel, the feeling of the time occupied by that stimulus^14^. While decades of research have built an understanding of the basic neuronal coding algorithms for stimulus features^4–10^, a mechanistic, causal understanding of the sense of time is still lacking. Here, we combine rat psychophysics with optogenetics to demonstrate that time perception may be treated with the language of sensory coding.

## Duration and intensity percepts interact

On each trial, rats compared two vibrissal vibrations (stimulus 1, stimulus 2; Fig. 1a). Vibrations were constructed by concatenating a sequence of speed values, sampled from a half Gaussian distribution^19^. A single vibration was defined by its intensity (*I*) in units of mean speed, and its duration (*T*). We trained two sets of rats. *Duration rats* had to compare the two stimuli according to their relative time spans (*T1*>*T2* or *T2*>*T1*). *Intensity rats* had to compare the two stimuli according to the analogous relation (*I1*>*I2* or *I2*>*I1*). The two groups received the same stimulus set (Extended Data Fig. 1), the only difference being the feature they were trained to extract – for duration rats, stimulus intensities were irrelevant to the task, while for intensity rats stimulus durations were irrelevant (Fig. 1a, gray and red arrows prior to choice).

**Figure 1.**
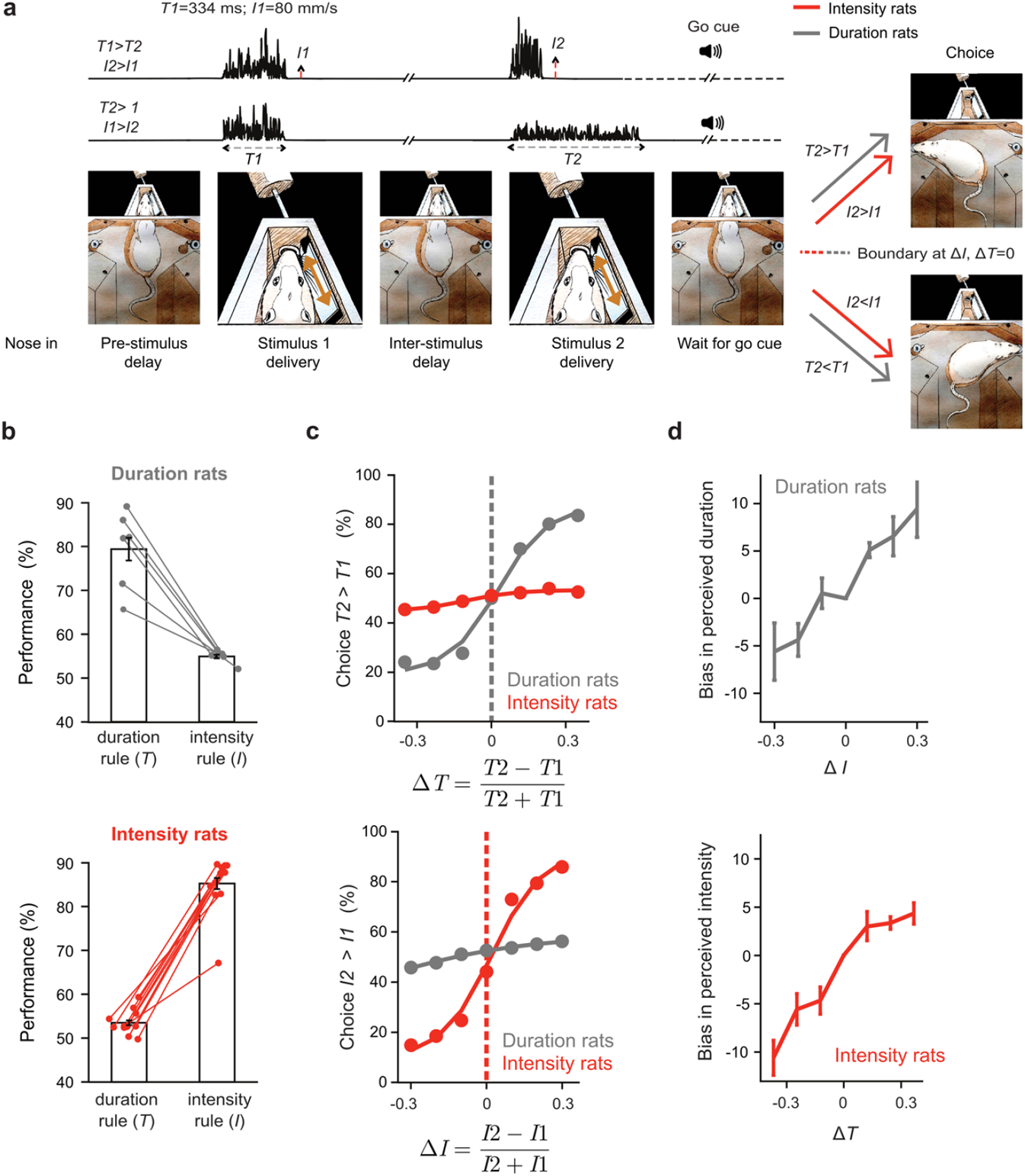
Interacting perception of duration and intensity. **a**, Rat enters the nose poke, bringing its right whiskers into contact with the plate. Following a pre-stimulus delay (0.5 s), stimulus 1 is delivered through plate motion. Stimulus 2 is presented after the inter-stimulus delay (2 s). Acoustic go cue prompts the rat to make a choice. **b**, Upper: performance of duration rats (n=6) based on duration and intensity rules. Lower: performance of intensity rats (n=11). Analysis of choices according to duration rule done on trials with largest Δ*T*; analysis of choices according to intensity rule done on trials with largest Δ*I*. **c**, Upper: Psychometric curves based on Δ*T*. Lower: Psychometric curves based on Δ*I*. **d**, Upper: Intensity-dependent bias in perceived duration. For a given Δ*I*, bias is the average of the percent of trials judged *T2*>*T1* across *ΔT* values. Lower: Duration-dependent bias in perceived intensity, computed in the analogous way. Bias measure details in Methods.

In the upper plot of Figure 1b, the left bar depicts the performance (78% correct) of duration rats when choices are analyzed according to relative stimulus durations. When the same choices are analyzed according to relative stimulus intensities (right bar), performance was above chance (54% correct). Intensity rats performed at 85% correct according to relative stimulus intensities (lower plot, right bar); when analyzed according to relative stimulus durations (left bar), performance was above chance (53% correct).

The psychometric curves of Figure 1c show choices according to graded stimulus differences. In the upper plot, choices in duration rats (gray) were governed by Δ*T* (normalized duration difference, defined as (*T2-T1*)/(*T2*+*T1*)) while choices in intensity rats (red) were weakly modulated by Δ*T*. In the lower plot, choices in intensity rats (red) were governed by Δ*I* (normalized intensity difference, defined as (*I2*–*I1*)/(*I2*+*I1*)) while choices in duration rats (gray) were weakly modulated by Δ*I*.

To quantify the bias caused by the irrelevant feature, intensity, on duration perception, we computed how the likelihood of judging *T2*>*T1* was affected by Δ*I* (Fig. 1d, upper plot; also see Methods). Similarly, we quantified the duration-dependent bias in perceived intensity by computing how the likelihood of judging *I2*>*I1* was affected by Δ*T* (Fig. 1d, lower plot).

## Optogenetic control of perception

Motivated by the interaction between perceived duration and perceived intensity (Fig. 1b-d), the remaining experiments test the hypothesis that vibrissal somatosensory cortex (vS1) forms the bases for rats’ judgment of both features; further analyses seek to specify the underlying neuronal code. ChR2(H134R) or eNpHR3.0 was expressed in left vS1 (Fig. 2a, left) and neuronal populations were accessed by movable microdrive arrays coupled with optic fibers (Fig. 2a, middle). If vS1 directly participates in the brain’s perceptual clock, optogenetic excitation or inhibition of vS1 (Fig. 2a, right) will affect the judgment of time. When blue light was applied in EYFP-ChR2(H134R)-expressing duration rats, optogenetic excitation during presentation of stimulus 2 caused a leftward shift of the psychometric curve, indicating an *overestimation* of that vibration’s duration. The opposite shift, rightward, was obtained with optogenetic excitation during stimulus 1 (Fig. 2b). When red light was applied in eNpHR3.0-expressing duration rats, optogenetic inhibition during the presentation of stimulus 2 or stimulus 1 caused *underestimation* of that vibration’s duration (Fig. 2c). In intensity rats expressing EYFP-ChR2(H134R), optogenetic excitation during presentation of stimulus 2 or stimulus 1 caused *overestimation* of that vibration’s intensity (Fig. 2d).

**Figure 2.**
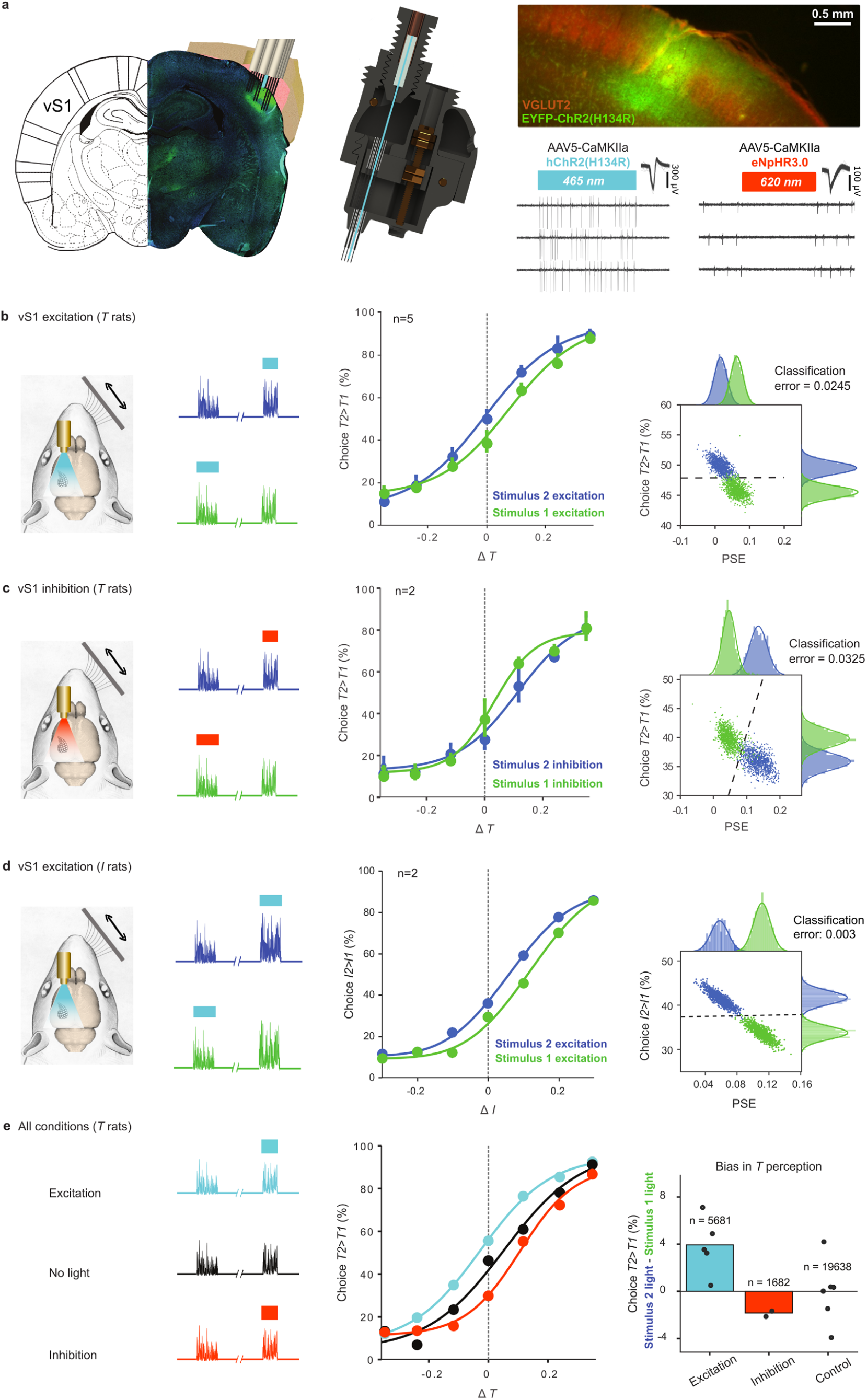
Optogenetic manipulation of time and intensity perception. **a**, Left: EYFP-ChR2(H134R)-injected brain with optical fiber surrounded by an array of electrodes. Middle: custom-built multisite drivable optrode array. Upper right: coronal section of the EYFP-ChR2(H134R) injection site (green) counterstained with Anti-VGlut2 primary antibody (red). Lower right: traces of two vS1 single-neurons. In the EYFP-ChR2-injected rat 465 nm illumination (blue bar, 500 ms) excites the neuron while in the eNpHR3.0-injected rat 620 nm illumination (red bar, 250 ms) inhibits the neuron. **b**, Left: optogenetic excitation of left vS1 during vibrissal stimulation (left vibrissae not shown). Middle: excitation during stimulus 2 (blue); excitation during stimulus 1 (green). Right: two signatures of the curve shift, percent of trials judged as *T2*>*T1* irrespective of stimulus duration (ordinate) and PSE, the point of subjective equality (abscissa), were measured with bootstrap resampling methods. A support vector machine classifier quantifies data separation by classification error. **c**, Same as **b**, but for optogenetic inhibition. **d**, Same as **b**, but for intensity rats. **e**, Left and middle: optogenetic excitation (blue) during stimulus 2 yielded an overestimation of *T2*, while optogenetic inhibition (red) during stimulus 2 yielded an underestimation of *T2*. Right: Bias in duration perception induced by optogenetic manipulation.

Blue light in the apparatus, applied with the same temporal alignment to the vibration, had no effect (Extended Data Fig. 2), indicating that behavioral biases were not due to visual cues. While causing a shift in psychometric curves, optogenetic stimulation did not alter the overall accuracy (Extended Data Fig. 2).

The effects of optogenetic stimulation (vS1 excitation, inhibition, and controls) are pooled in Figure 2e. The right panel illustrates the bias in perceived duration, measured by the method of Figure 1d. Notwithstanding individual differences in the magnitude of effect, rats showed a significant bias towards judging a vibration as having extended duration (blue) or compressed duration (red) when accompanied by vS1 excitation and inhibition, respectively.

## Time coding

Through what physiological mechanisms does the neuronal firing within vS1 give rise to the percept of the passage of time? Figure 3a illustrates two example neurons from EYFP-ChR2(H134R)-expressing rats, representative of the population’s heterogeneity in responsiveness to vibrissal stimulation and optogenetic excitation^20^. The neuron in the left plots responded weakly to vibrissal stimulation, showing only an onset transient. The same neuron was robustly excited by blue light. The neuron in the right plots gave a strong, non-adapting response to vibrissal stimulation, but was not excited by blue light. Sensory responses, with and without optogenetic excitation, are shown in Figure 3b. Color indicates the deviation of firing rate from the baseline level (z-score). In the upper plot, neurons are ordered by response magnitude to stimulus 2, which was accompanied by optogenetic excitation. In the lower plot, the neurons are ordered by response magnitude to stimulus 1, which was accompanied by optogenetic excitation. The population’s response to vibrissal stimulation alone and vibrissal-plus-optogenetic excitation can be seen by comparing the upper and lower plots (quantification in Extended Data Figs. 3, 4).

**Figure 3.**
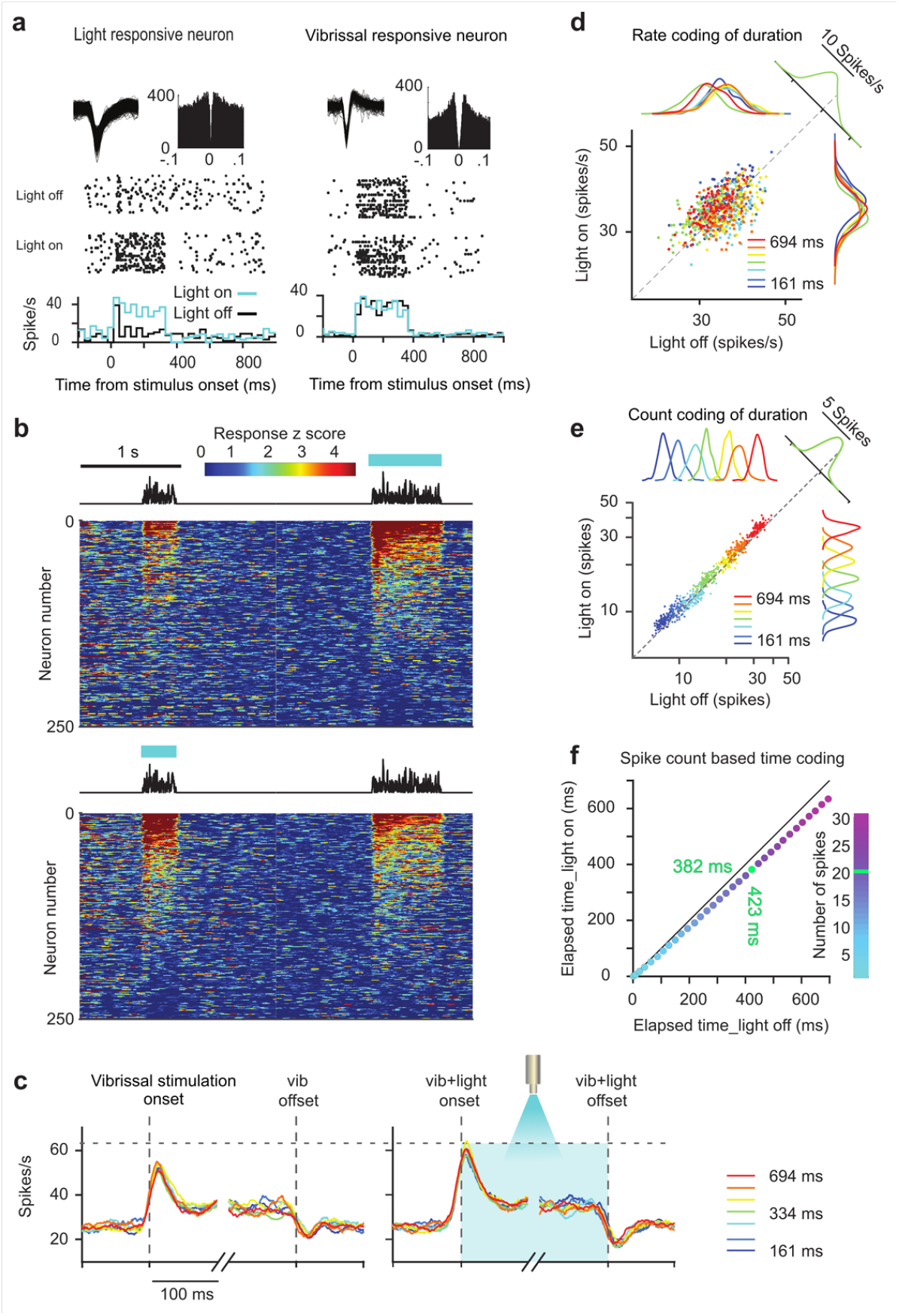
vS1 coding of duration. **a**, Example vS1 neurons. Upper: spike waveforms and spike time autocorrelogram. Middle: raster plot on randomly selected light-off and light-on trials (334 ms stimulus duration). Lower: both neurons’ firing rates in non-overlapping 20 ms bins grouped by light-on (light blue) and light-off (black) trials. Stimulus duration was 334 ms. **b**, Normalized response of all neurons on trials in which stimulus 1 was 334 ms and stimulus 2 was 694 ms. **c**, Average response of all neurons during the first and final 100 ms of stimulus 2 presentation without (left) and with (right) blue light. Color denotes stimulus duration. Light onset and offset (right) matched vibrissal onset and offset. Neuronal responses appear to rise before stimulus onset due to temporal “leakage,” as PSTH values are derived by a centered 20 ms-sliding window. **d**, Each dot shows population mean firing rate (bootstrap resampling) colored by stimulus duration. Distributions for each duration are shown as marginals. Dashed diagonal denotes equal firing rate for light-on and light-off. For 334 ms duration, points are projected parallel to the diagonal to give the green histogram. **e**, Same as **d**, but for spike count summated across the entire stimulus presentation. **f**, Points depict the elapsed time required to reach a given spike count (color scale) during stimuli with the light off (abscissa) and light on (ordinate). The points’ positions below the solid diagonal indicate that a given spike count was reached earlier with optogenetic excitation. Data obtained by population response resampling.

The temporal profile of vS1 sensory responses was conserved under optogenetic excitation^21^. Figure 3c, left, shows the population peristimulus time histogram (PSTH) in EYFP-ChR2(H134R)-expressing rats in the absence of optogenetic excitation. To compare across different durations, the first and final 100ms are shown. After an early peak, firing rate remained stable until offset. The right panel shows the PSTHs of the same population under optogenetic excitation, revealing an evenly distributed boost in the vibrissae-evoked response.

Two coding regimes fundamental to sensory processing are firing rate (number of spikes per fixed time window) and spike count (the summated number of spikes, from stimulus onset to offset)^4–7^. If firing rate functions as an explicit representation of stimulus duration within vS1, it must vary systematically in relation to the passage of time. Figure 3d examines the firing rate of the entire recorded vS1 population during the final 100 ms of stimulus presentation, with light-off (abscissa) and light-on (ordinate). Points were obtained by bootstrap resampling and colors denote stimulus duration. Projection along the diagonal reveals the overall effect of optogenetic excitation – a leftward shift towards higher firing rate (depicted only for the 334 ms duration, green). However, when the points are projected laterally and vertically, the marginal distributions for each duration are fully overlapping. Thus, firing rate at the end of the stimulus, although boosted by the optogenetic intervention, does not robustly encode duration.

As an alternative, Figure 3e examines the spike count as the possible basis for a duration code. Following the format of the preceding panel, counts from vibrissal-stimulus onset to offset, with light-off (abscissa) and light-on (ordinate) were computed, and the projection along the diagonal reveals a leftward shift towards greater spike count under optogenetic excitation (depicted only for the 334 ms duration). Differently from the rate code, when points are projected laterally and vertically, marginal distributions separate according to stimulus duration. Thus, summated spike count could provide a downstream integrator with an input that robustly encodes duration, and is consistent with the dilation of perceived time generated by optogenetic excitation.

If a downstream integrator were to use a simple spike count code, what would be the magnitude of the optogenetic effect? We plotted how much time from stimulus onset must have passed in trials with light-off and light-on to reach any selected count of spikes (Fig. 3f). For instance, when 20 spikes (data point in green) have been integrated, 423 ms would have passed on trials with light-off, but just 382 ms on trials with light-on; optogenetic excitation would cause the 382 ms-stimulus to be perceived as 41 ms longer.

## Model of coding mechanisms

Figure 3e-f posits the perceptual effects of *linear* summation of vS1 spikes, but the dynamics of downstream integration are likely to be *non-linear*^5,14^. Therefore, we grounded the search for the mechanisms of time perception in a 3-stage model encompassing non-linear integration. In stage 1, vibrissal drive and optogenetic drive evoke currents in vS1 neurons, leading to spiking through a linear-nonlinear Poisson (LNP) process (Fig. 4a, left). After finding the parameters that produce simulated spike trains mimicking the original spike trains (stimulus-dependence, variability and diversity), we recombine these currents in a Gaussian Mixture Model^22^ to create a large pool of simulated vS1 neurons (Extended Data Fig. 5). In stage 2, the accumulated quantity (*ϒ*) in a leaky integrator (LI) downstream to vS1 is taken as the duration percept. As the integrator summates incoming spikes, input continuously leaks out by some proportion (*ϒ*/*τ*) (Fig. 4a, upper right). In stage 3, the values of *ϒ* at the conclusion of stimuli 1 and 2 are taken as the explicit readouts of duration and their comparison predicts the rat’s actual choice (Fig. 4a, lower right).

**Figure 4.**
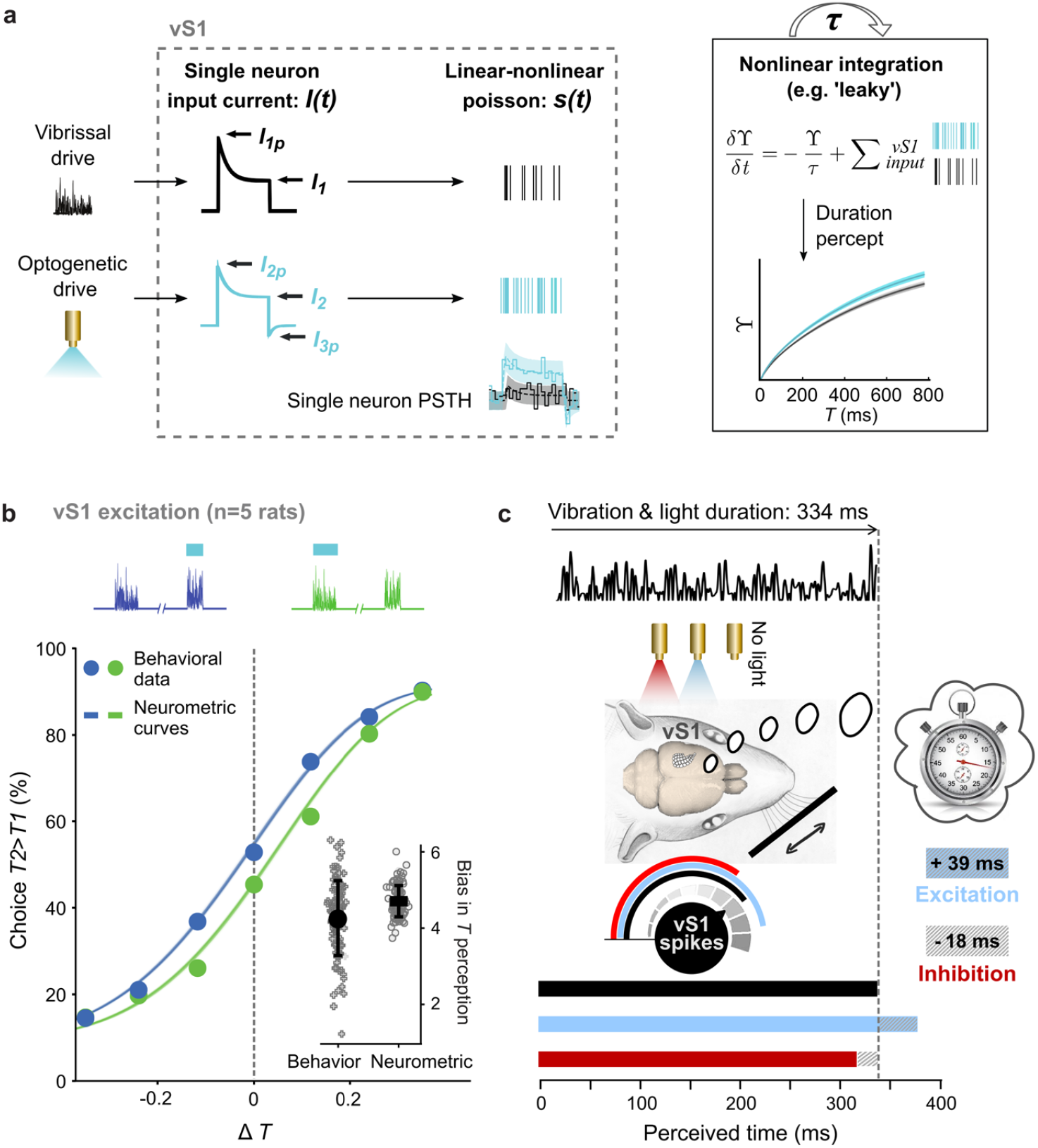
Model of non-linear integration to generate time percept. **a**, Left box: stimulus-dependent input currents that follow characteristic dynamics for vibrissal and optogenetic drive, converge on vS1 neurons, giving rise to spike trains with Poisson statistics. Example vS1 PSTH is shown on the lower right corner; binned neuronal activity (solid line) in response to a vibrissal stimulus (black), as well as with optogenetic excitation (light blue), are simulated (dashed line with confidence intervals) by fitting the respective input currents and feeding them to an I/F. Current and I/F parameters in Methods. We simulated a 5000-neuron population based on the distribution of fitting parameters of the entire population of recorded neurons. Right box: the LI receives input spike trains from the simulated vS1 population under conditions including vibrissal (black spike train) and vibrissal plus optogenetic input (blue spike train). The integrator’s accumulated quantity is governed by the differential equation. Reading out the generated vS1 neuronal population activity with this LI, we predict the perceptual shift created by optogenetically increasing firing rate in vS1. **b**, Psychometric curves of the neurometric model (points). Inset: comparison of the perceptual shifts between behavioral data and generated neurometric curves for resampled (100x) neuronal and behavioral data. Bias is quantified by the measure introduced in Figure 1d. **c**, vS1 role in compressing or dilating perceived time by its sensory drive; optogenetic manipulation slows or speeds the perceptual “stopwatch.” Changes in perceived time derived from the shift in the point of subjective equality (PSE) in the averaged behavioral data.

This model yields the neurometric curves (solid lines) of Figure 4b, which overlie the observed psychometric data (points), indicating that the model offers a physiologically plausible framework for how vibrissal drive and optogenetic excitation of sensory cortex generate perceived duration. The scatter plots of Figure 4b (inset) show the optogenetic excitation-induced bias in perception, in behavioral data and modelled neurometric output.

The distance between the blue and green curves (Fig. 4b) allows estimation of the direct perceptual effect of vS1 intervention (Fig. 4c). A vibrissal stimulus of actual duration 334 ms, absent any direct intervention in sensory cortex, will be perceived as having a veridical duration of 334 ms (black bar). That same stimulus, when accompanied by vS1 excitation (blue light) will be perceived (on average) as having a duration of 372 ms, an optogenetic-derived perceptual dilation of 39 ms. When accompanied by vS1 inhibition (red light), that stimulus will be perceived (on average) as having a duration of 316 ms, an optogenetic-derived perceptual compression of 18 ms. The empirical observations, coupled with the physiological model for vS1 and downstream integration, offer a detailed picture for how the perceptual clock embodies sensory coding in the cortex.

## Discussion

Applying targeted and controlled optogenetic manipulation of vS1 in one set of rats performing tactile duration discrimination and in another set of rats performing tactile intensity discrimination, this study reveals parallel generation of two percepts, with primary sensory cortex common to both networks. While different mechanisms may be at work in processing the “empty” interval between two discrete events^23–25^, time perception accompanying an ongoing stimulus stream arises within the sensory representation of touch. The contribution of the coding of stimulus features to sense of time distances our findings from dedicated pacemaker-accumulator operation models^11–13^, where the accumulator receives onset/offset triggers but is insensitive to the neuronal coding of the stimulus. In existing models where the time percept is constructed from sensory drive^14,15^, no causal link between sensory cortex and the final percept has yet been established.

Embodied within a network extending to the sensory cortex, time perception now becomes amenable to the tools previously restricted to quantifying the representation of stimulus features – tools such as spike counts, firing rates and temporal patterns^4–10^. In short, time perception can be treated through the computational language of sensory coding. One immediate insight from this treatment is that processing likely involves integration of sensory cortical input with long integration time constants, as are found in frontal cortical regions^5^.

A crucial component of the present model, the accumulation of sensory drive by a downstream integrator, appears to apply to humans^14,16^. The generality of the model raises the prospect that anomalous sensory coding mechanisms may be one contributing factor in the time misperception at the core of multiple psychiatric disorders^17,18^.

## Methods

### Rat subjects

All protocols conformed to international norms and were approved by the Ethics Committee of SISSA and by the Italian Health Ministry (license numbers 569/2015-PR and 570/2015-PR). 20 male Wistar rats (Harlan Laboratories, San Pietro Al Natisone) were caged in pairs and maintained on a 14/10-hour light/dark cycle. They were trained and handled on a daily basis and provided with daily environmental and social enrichment. To promote motivation in the behavioral task, rats were water-restricted for approximately 20 hours prior to training or testing sessions; access to food in the cage was continuous. They were tested each weekday in sessions of about 1 hour.

### Behavioral task

To initiate a trial, the rat entered the nose poke, placing its whiskers in contact with a plate connected to a shaker motor (type 4808; Brüel & Kjær see^19^). It then received two vibrissal stimuli separated by a delay. Stimuli were noisy vibrations, constructed by stringing together over time a sequence of plate velocity values (motion along the axis of the rod connecting the plate to the motor). Velocities were sampled from a Gaussian distribution with 0 mean and standard deviation ranging from 25 to 148 mm/s. The speed distribution (absolute values of velocity) was a half-normal (folded) distribution whose mean was equivalent to the standard deviation of the underlying Gaussian multiplied by 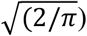. We refer to mean speed as intensity (*I*). Vibration duration was denoted *T*. Durations varied from 112 to 1000 ms (see Extended Data Fig. 1). The differences between the two stimuli making up one trial are expressed by two indices, normalized intensity difference (Δ*I*) and normalized time difference (Δ*T*):

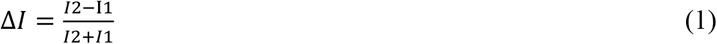

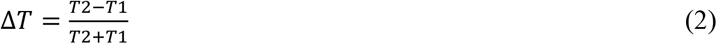

where *I1* and *I2* are the intensities, and *T1* and *T2* are the durations, of stimuli 1 and 2, respectively.

*Duration rats* were trained and tested using a rule where reward location was determined by the sign of Δ*T*, with Δ*I* irrelevant. *Intensity rats* were trained and tested using a rule where reward location was determined by the sign of Δ*I*, with Δ*T* irrelevant.

In test sessions, each stimulus combination assumed duration and intensity values from the stimulus generalization matrix (see Extended Data Fig. 1). Thus, rats received a random combination of 10 intensity pairs (*I1, I2*) x 10 duration pairs (*T1, T2*) during each training session. Moreover, in each trial, the relevant and irrelevant features could be congruent (*I2*>*I1, T2*>*T1* or *I1*>*I2, T1*>*T2*) or incongruent (*I2<I1, T2*>*T1* or *I1*<*I2, T1*>*T2*). This randomness in congruence required the rat to act upon only the relevant feature in order to perform above chance. To obtain psychometric curves for duration, *T1* took a fixed value of 334 ms while *T2* spanned seven possible durations, giving a range of Δ*T* from -0.35 to 0.35. The same design was used to obtain psychometric curves for intensity, with *I1* fixed at a 65 mm/s, while *I2* spanned seven possible values; Δ*I* ranged from -0.3 to 0.3.

### Targeted virus injections in vS1

After rats reached stable behavioral performance, they were anesthetized with 2-2.5% Isoflurane in 100% oxygen delivered through a customized plastic snout mask. Target regions were accessed by craniotomy, using standard stereotaxic technique. The vasculature visible on the brain surface was used as a reference for cortical maps^26^. Photos of the brain surface of vS1 were made with a 5x Zeiss microscope connected to a webcam and further used to document electrode insertion and injection sites. Single tungsten electrodes (100-500 kΩ impedance; FHC) were inserted to a depth of ∼750 µm and the whisker constituting the neuronal population’s strongest input was assessed by stimulation with a hand-held probe. Neuronal populations with receptive fields on whisker rows C-D and columns 4-6 were targeted. AAV5-CaMKIIa-hChR2(H134R)-EYFP or AAV5-CaMKIIa-eNpHR3.0-EYFP (UNC vector core) was prepared by standard procedures^27^. A 10 µl Hamilton syringe was filled with 4-6 µl of virus solution (stained with Fast Green FCF). Injections of 0.5 µl of virus solution were made at depths of 800 and 1600 µm in 3-4 barrel-columns with identified receptive fields. The skull opening was conserved with custom-made cylindrical implanted cranial windows. Four to six screws were fixed in the skull as support for dental cement. Two screws served as reference and ground and were connected via a silver wire to a 2-pin connector that was embedded in dental cement. At the conclusion of the operation, rats were treated with antibiotic (Baytril; 5 mg/kg; i.p.), analgesic (Rimadyl; 2.5 mg/kg, i.m.), atropine (ATI; 2 mg/kg, s.c.) and with sterile saline to rehydrate (5 ml, s.c.). A local antibiotic ointment was applied around the cutaneous wound to improve the healing. Tissue was washed regularly and treated with antibiotics in the weeks after surgery through the cylindrical implanted cranial windows. During the recovery period, rats had unlimited access to water and food.

### Implantation of opto-electric microdrive

In a second operation 3-4 weeks after virus injection, the opto-electric microdrive was implanted through the cranial window. Injection sites were identified by vasculature landmarks and mapping from the previous surgery. For chronic electrophysiological recording concomitant with optogenetic stimulation, we collaborated with CyNexo to design an opto-electric microdrive (aoDrive, https://www.cynexo.com/portfolio/neural-drives/, see also Fig. 2a, middle). Each drive incorporates up to 15 single FHC tungsten electrodes and an optic fiber (Ø: 230 µm, NA: 0.67, Plexon). Electrodes and fiber can be independently moved in depth with a total range of 4.5 mm. Electrodes were lowered until neuronal responses to light delivery (PlexBright LED, *Blue*: 465 nm, or *Orange*: 620 nm for inhibition, Plexon) were observed. Subsequently, microdrives and TDT connectors were embedded in dental cement. The opto-electric microdrive provided signals for 3-6 months after the surgery. Electrophysiological recording and optogenetic stimulation in the behaving animal began 7-10 days following the second implantation surgery.

### vS1 recordings and optogenetic behavioral experiments

Extracellular activity was pre-amplified, filtered and digitized using the digital TDT recording system (Tucker David Technologies) along with task-relevant data, such as position sensors and light/motor stimulation signals to synchronize external events with physiological recordings. Signals were sorted into single and multi-unit neuronal clusters, as verified through standard indices using UltraMegaSort2000^28^. The headstage and optic fiber patch cables (custom made, Ø: 230 µm, NA: 0.67) were connected to the implant and the cables were held by a rubber band to limit weight on the implant. Light output intensity (>=10 mW) from the tip of the patch cable was measured (Thorlabs) weekly to ensure stable optogenetic excitation/inhibition effect. Light delivery (465 nm) for the optogenetic excitation and external light experiment was 60 ms delayed to the onset of vibrissal stimulation to ensure that the light evoked responses did not prolong the elapsed time of vS1 excitation. Offset time of light delivery and vibrissal stimulation was identical. Light delivery (620 nm) for the optogenetic inhibition experiment was slightly adapted in order to account for the biophysical mechanisms of eNpHR3.0 (see^29^): (1) light was initiated 50 ms preceding onset of vibrissal stimulation to reach an effective hyperpolarization, accounting for the slower time constants of eNpHR3.0 as compared to Chr2(H1340). (2) Light was dimmed with an offset ramp, starting 200 ms before and ending at 0 ms with respect to termination of vibrissal stimulation; this protocol minimized rebound activation.

### Histological examination

At the conclusion of the study, electrolytic lesions were made around the tips of the electrodes to mark the recording sites. To identify Opsin-expressing neurons, we counterstained with blue-fluorescent Nissl stain (NeuroTrace 435/455, ThermoFisher) to visualize cortical layers, and performed antibody staining (AntiVGLUT2, Synaptic Systems) to discern the barrels in layer IV. Whole coronal slice (50 µm thickness) images were taken with confocal microscope (4x, Nikon).

### Analysis of behavioral data

To generate duration psychometric curves, we used trials in which *T1* was 334 ms while *T2* ranged from 161 ms to 694 ms. For intensity psychometric curves, we used trials in which *I1* was 64 mm/s while *I2* ranged from 34 mm/s to 119 mm/s (Extended Data Fig. 1). The rat’s choice (proportion of trials in which the stimulus 2 was judged as more intense or longer in duration than stimulus 1) for each stimulus pair was then plotted. A four-parameter logistic function was fit to the psychometric data using the nonlinear least-squares fit in MATLAB (MathWorks, Natick, MA), as follows. The psychometric curve for duration was given by

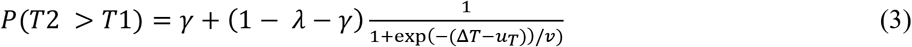

and for intensity the curve was given by

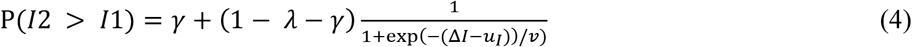

where Δ*T* and Δ*I* are the normalized stimulus differences, γ is the lower asymptote, λ is the upper asymptote, *1/ ν* is the maximum slope of the curve and *u*_*T*_ and *u*_*I*_ are Δ*T* and Δ*I* at the curve’s inflection point, for the duration and intensity curves, respectively. Since observed experimental data are expressed as choice (%), the proportion of trials in Eqs (3)-(4) was multiplied by 100 for purposes of illustration.

The raw data (choices for each value of Δ*T* and Δ*I*) as well as the fit parameters were then used as measures of acuity and bias. Bias in perceived duration was measured by sorting the trials according to Δ*I* and then, with the Δ*T* values pooled, averaging the percent judged *T2*>*T1* for each value of Δ*I*. This isolates the effect of Δ*I* on perceived duration. For illustration (Fig. 1d), the above bias measure was normalized by subtracting the choice data for Δ*I* = 0 (where *I* does not exert a bias). Easy trials (e.g., Δ*T* = + / -0.35) were excluded from this analysis as the irrelevant feature (Δ*I*) no longer exerts an effect.

### Resampling and bias statistics

To quantify the effect of optogenetic manipulation of vS1 on duration and intensity perception, we resampled the original data set to create 1000 sets of statistically comparable data. Each set of resampled responses was parametrized by fitting the logistic function of Equation (3). A support vector machine (SVM) classifier (MATLAB, *fitcsvm* function) was used to quantify the linear separation between data points with and without optogenetic intervention, making use of 10-fold cross validation to measure the classification error. Specifically, the data were partitioned into 10 random sets. Then, 9 of these were used to train an SVM classifier and the remaining set served as a test. This procedure was repeated 10 times and the statistics for each repetition were combined, giving the rightmost plots of Figure 1b-d.

### Neuronal data

Spike trains were aligned to the stimulus onset or else offset, depending on the aim of the analysis. In Figure 3b, individual neurons’ response was generated by plotting the average firing rate over trials, shifting in 1 ms steps. To reduce the effect of noisy fluctuations, a centered 40 ms sliding window was used. Firing rate was then z score– transformed by subtracting each neuron’s spontaneous activity rate (measured from 800 ms before stimulus onset up to the stimulus onset) from response rate per time bin during stimulus presentation. The outcome was divided by spontaneous activity variance.

Population PSTHs (Fig. 3c) were generated by plotting the average population response shifting in 1 ms steps for each stimulus duration. To reduce the effect of noisy fluctuations, a centered 20 ms sliding window was used. This led to the appearance of a rise in firing rate just before stimulus onset and a decrease in firing rate just prior to stimulus offset.

### Model for vS1 activity

The analysis was built on the spiking activity of 240 vS1 units recorded in 5 rats over the full range of vibrissal stimulation and optogenetic excitation conditions. To permit more robust neurometric measures, we modeled a larger data set replicating the properties of actual recordings by means of a Gaussian Mixture Model (GMM)^22^. The key observations justifying this form of model are that functional properties are diverse across neurons and the responses of single neurons are variable across trials. The methodology is presented in two steps: (1) constructing a parametric model for single neuron variability and fitting it to each recorded unit, and (2) Gaussian mixture model to reproduce vS1 population diversity.

(1) The parametric model for neuron variability is based on the following observations: (i) the response of a neuron varied across trials notwithstanding constant stimulus conditions, (ii) in the absence of stimulation, vS1 neurons showed ongoing activity, (iii) when the vibrissal stimulus was presented, many vS1 neurons rapidly increased their firing rate and then adapted to steady state, (iv) simultaneous optogenetic and vibrissal stimulation commonly led to a higher firing rate but did not alter the temporal profile of response. However, firing rate sometimes dropped after the light was turned off (a phenomenon sometimes referred to as post-stimulation suppression^30^.

Given the recorded units’ observed spike variability, we assume that the activity of the *i*^th^ unit, *k*^(*i*)^, in time bin *t*, follows a Poisson distribution.

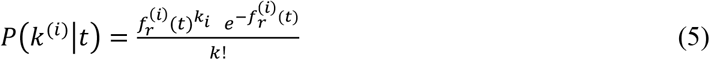

where *f*_*r*_^(*i*)^(*t*) is the unit’s firing rate. We write a parametric model for the underlying rate of the Poisson distribution and infer its parameter values based on the measured PSTH of each recorded unit. This spike generation model assumes that the *i*^th^ unit receives a constant background current *I*_*0*_^(*i*)^. We assume that when the vibrissal stimulus is turned on, it elicits a mechanoreceptor-derived current of the following form

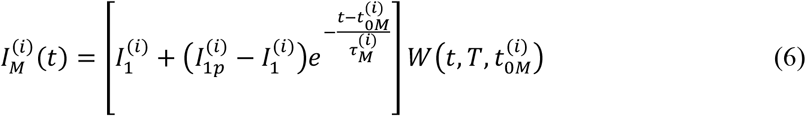

Where *I*_1*p*_^(*i*)^ is the peak input current that follows immediately after stimulus onset, *I*_1_^(*i*)^ is the steady state current, *t*_0M_^(*i*)^ is the onset of the mechanoreceptor-elicited current, and *τ*_*M*_^(*i*)^ is the decay time constant from *I*_1*p*_^(*i*)^ to *I*_1_^(*i*)^. *W(t, T, t*_0M_^(*i*)^) is a window function that determines when the stimulus evokes a current that is non-zero:

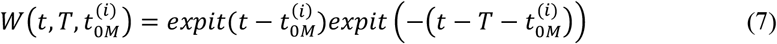

where *expit*(*x*) = 1/(1 + *exp*(−*x*)).

When the optogenetic excitation is turned on, it elicits a current of the following form:

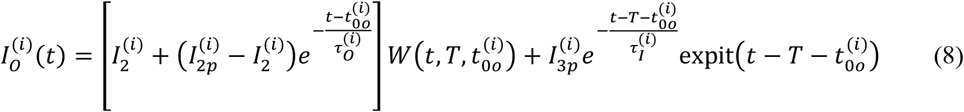

where 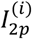 is the peak input current that follows immediately after light onset, 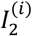 is the steady state current, 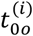 is the time of onset of the current, 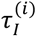 is the decay time constant from 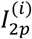 to 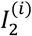, and 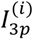 is the activation current that follows light offset. 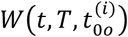 is the same windowing function as in the mechanically elicited current.

The total input current received by the *i*^th^ unit is

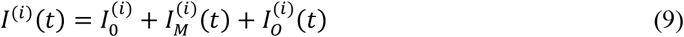

Input current to output firing rate curve (I/F curve) is modeled as a generalized sigmoid^31^:

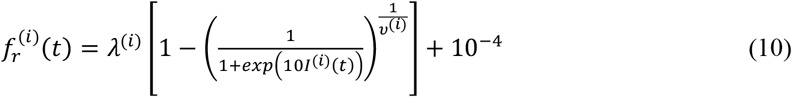

Where *λ*^(*i*)^ denotes the maximum firing rate and *ν*^(*i*)^ represents the non-linear scale of the generalized sigmoid curve. The current *I*^(*i*)^(*t*) is multiplied by a constant scaling factor 10, to improve the fit stability due to the different scales of the many parameters involved. The added 10^−4^ constant helps to stabilize the fit in very low-firing neurons.

We used the pymc3 package^32,33^ to infer the 12 parameters given the observed PSTH of the neurons in 10 ms wide time bins. The model output accurately fitted the mean firing rate and the variability in the spike trains for the 240 recorded units (see S4).

(2) The Gaussian mixture model exploits the diverse properties of individual vS1 neurons to make estimates of large populations on the basis of limited quantities of recordings. The reasoning is that a recorded neuron, unit *i*, is a randomly sampled member of a broader group of vS1 neurons with similar properties (i.e., similar parameters according to the single neuron variability model described above). We label this group *g*_*i*_. Unit *i* emits spikes with a Poisson probability distribution (Equation (10)).

In detail, we first assume that vS1 is made up by a mixture of *G* qualitatively different classes of neurons, where classes are defined by the parameter values of the single neuron model. Each class can be represented as a Multivariate Gaussian in a subspace *A* of the model’s parameter space, where A is determined by *I*_0_, *I*_1_, *I*_1*p*_, *I*_2_, *I*_2*p*_, *I*_3*p*_, *λ* and *ν*. The parameters, *τ*_*M*_, *τ*_*O*_ and *τ*_*I*_ showed only minimal variations among neurons, making it likely that they are population-specific and not neuron-specific; thus we chose to fix their values to the median of the fit for all neurons: *τ*_*M*_ = 48 ms, *τ*_*O*_ = 49 ms and *τ*_*I*_ = 28 ms. The parameters *t*_0*M*_ and *t*_0*o*_ are defined by the experimenter and therefore kept constant.

Class membership is derived from a Dirichlet process^34^. The full generative process underlying real neuronal data can be written as

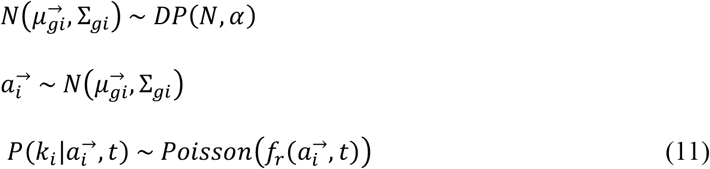

where the *i* subindex corresponds to the recorded unit, the *g*_*i*_ subindex represents the group to which the *i*^th^ unit belongs to, 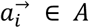 is the vector of parameters 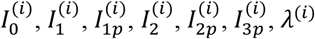 and *ν*^(*i*)^, *DP* is a Dirichlet process, *α* is the concentration parameter, and 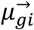 and Σ_gi_ are, respectively the g_i_^th^ class mean and covariance.

We infer the mixture weights, 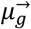 and Σ_*g*_ by using the expected values of the independently inferred parameters from the single neuron variability model as the observed 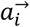 vectors for each unit. We then used scikit-learn’s^35^ built-in `BayesianMixtureModel` class to infer the suitable class proportions, and each class’s 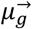 and *Σ* _*g*_. We set the *α* concentration hyperprior to 10^−6^ to favor assigning significant weight to a larger number of classes, but also set *G* = 5 to prevent excessive granularity.

This generative process and the Bayesian Mixture Model allow us to model a 5,000-unit vS1 population. The resulting spike trains are statistically consistent with observed single neuron variability and diversity in vibrissal stimulation and optogenetic excitation response. We take the full set of modeled neurons to be the drive *f*_*vS1*_*(t)* for any given stimulation condition, as

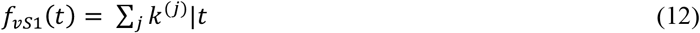

### Model for perceived stimulus duration

The perceived stimulus duration, *ϒ*, is modeled by the leaky integrator differential equation

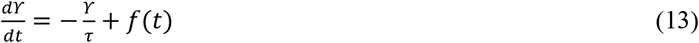

where *τ* is the leaky integrator time constant and *f(t)* is the external drive. The drive is written as

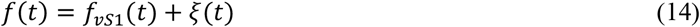

where *f*_*vS1*_(t) is neuronal activity in vS1, including vibrissal- or optogenetic -stimulation evoked responses and *ξ*(t) is ongoing firing unrelated to vibrissal or optogenetic stimulation. We approximate *ξ* as a Gaussian stochastic variable with mean *µ*_*b*_ and variance *σ*^*2*^_*b*_.

Leaky integration is not specified as a unique physiological process; rather, it represents the dynamics governing the percept (*ϒ*) in a manner that quantitatively accounts for the rat’s judgments as a function of combined vibrissal stimulation and optogenetic excitation evoked responses.

### Model for choice

Given the leaky integrator dynamics of *ϒ*(*t*), the model for *f*(*t*), and the approximation of the Poisson variability in *f*_*vS*1_ (*t*), we can solve the stochastic differential equation (13) as shown previously^14,36,37^). *ϒ(t*) follows a Gaussian distribution and its expected value and variance are equal to

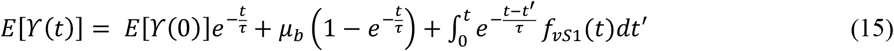

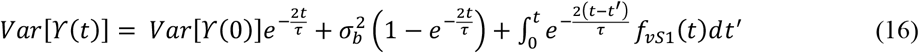

During stimulus delivery, the rat’s percept of elapsed time evolves through equation (13). The percept of total stimulus duration is given by *ϒ* at the time of stimulus offset (*ϒ*(*T*)). We then compute the probability distribution for each stimulus duration, *ϒ*(*T*1) and *ϒ*(*T*2), in the delayed comparison task. This gives the probability that *ϒ*(*T*2) is greater than *ϒ*(*T*1) as

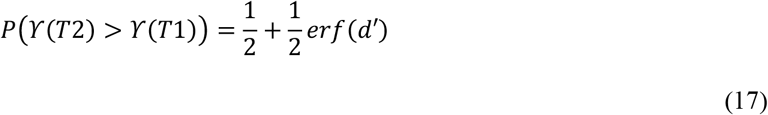

where

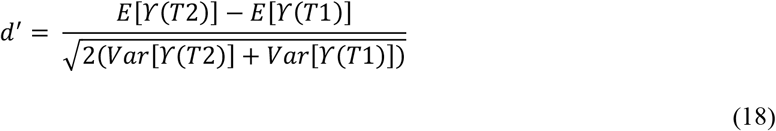

We can assume two types of trials^38^ – those in which the rat encoded *ϒ*(*T*2) > ϒ(*T*1) and used the two representations to make a choice (“attended” trials) and those in which choice was unrelated to the evoked sensory representations (“lapse” trials). We assume that in attended trials the rat judged *T2*>*T1* whenever *ϒ*(*T*2) > ϒ(*T*1); in lapse trials, the rat chose at random according to some choice bias probability *b*_*L*_. The probability that the rat judged *T2* > *T1* is then

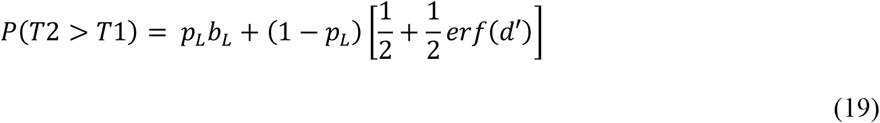

where *p*_*L*_ is the probability of a lapse trial.

### Fit of the behavioral psychometric data

We constructed 100 independent population proxies for vS1 firing, each made up of 5,000 neurons using the model described in (10). We then fit a common *τ, µ*_*b*_, 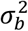, *p*_*L*_ and *b*_*L*_ across all 100 populations using a maximum likelihood estimate based on equation (19) with L2 parameter regularization for *τ*. The weight of the regularization was set to 0.01 and its center was placed at 600 ms. The resulting parameter values are listed in table (Extended Data, Tab. 1).

Using the resulting parameters for the neuronal population proxies, we computed the model’s predicted psychometric curves (Fig 4B), and the behavioral bias (overall predicted probability of choosing *T2* > *T1*, Fig 4B inset).

## Acknowledgments

We thank A. Toso, N. Nikbakht and J. Nicholls for helpful discussions and valuable insights and F. Manzino (CyNexo) and S. Parusso (CyNexo) for their invaluable technical assistance. A. Toso provided helpful comments on the manuscript. S. Sorella and B. Trovo assisted in animal training and behavioral data acquisition. We kindly acknowledge M. Riggi and M. Grandolfo for helpful comments on histology and imaging. M. Mahn, A. Akrami, J.-J. Sun are kindly acknowledged for helpful comments on optogenetics methodology. This work was supported by Human Frontier Science Program, project: RGP0015/2013 (MED), European Research Council advanced grant CONCEPT, project: 294498 (MED), European Union FET grant CORONET, project: 269459 (MED), Italian Ministry of Education, Universities and Research grant HANDBOT, project: GA 280778 (MED).

## Author contributions

A.F., S.R. and M.E.D. conceptualized the study and designed the experiments. S.R., M.G. and A.F. were designing and setting up the optogenetic methodology. L.P., S.R., A.F. and M.E.D. were conceptualizing, scripting and interpreting the computational model. S.R., A.F. and F.P. performed the experiments. A.F., S.R., L.P., M.E.D. performed the formal analysis. A.F., S.R. and M.E.D. performed data curation. M.E.D., S.R., A.F., L.P. and F.P. wrote the original draft and M.E.D., S.R., A.F. reviewed and edited the manuscript. M.E.D performed the funding acquisition.

## Data availability

The data that support the findings of this study are available from the corresponding author upon reasonable request.

## Code availability

The code for all analyses in this study is available from the corresponding author upon reasonable request.

**Extended Data Figure 1:**
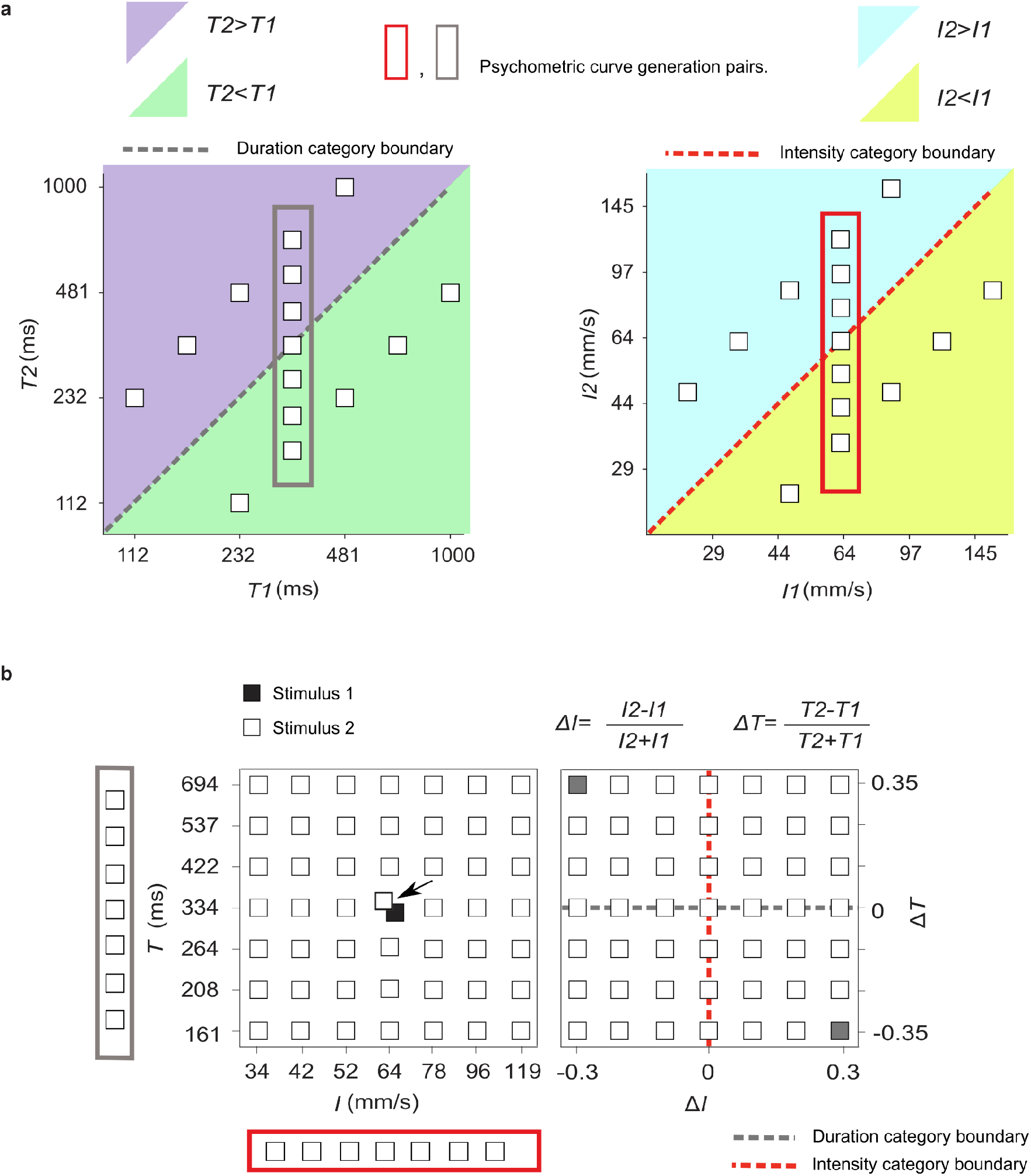
Stimulus generalization matrix. **a**, The entire stimulus set was used for both duration and intensity rats. *T1* and *T2* duration combinations are represented by blank squares (left panel). The diagonal gray dashed line indicates the category boundary (*T2*>*T1* vs *T2*<*T1*). The right panel accordingly displays possible *I1* and *I2* values that are relevant to the intensity task. For both, duration and intensity rats, all possible combinations of [*T1, T2*] and [*I2, I1*] were presented randomly across trials. The [*T1, T2*] and [*I2, I1*] combinations inside the gray and red rectangle are trials that were used to generate psychometric curves. **b**, Stimulus combinations that were used to generate psychometric curves in order to assess bias and acuity. Left panel: trials in which *T2* and *I2* (empty squares) varied in small steps, while *T1* and *I1* (central filled square) were fixed (334 ms, 64 mm/s). Right panel illustrates the normalized duration and intensity differences, corresponding to the stimulus pairs in the left panel. Duration rats must base their choice on Δ*T* values (above and below the gray dotted line), while intensity rats must choose based on Δ*I* values (left and right side of the red dotted line). Gray filled squares represent stimulus pairs illustrated in Figure 1A example stimulus traces.

**Extended Data Figure 2:**
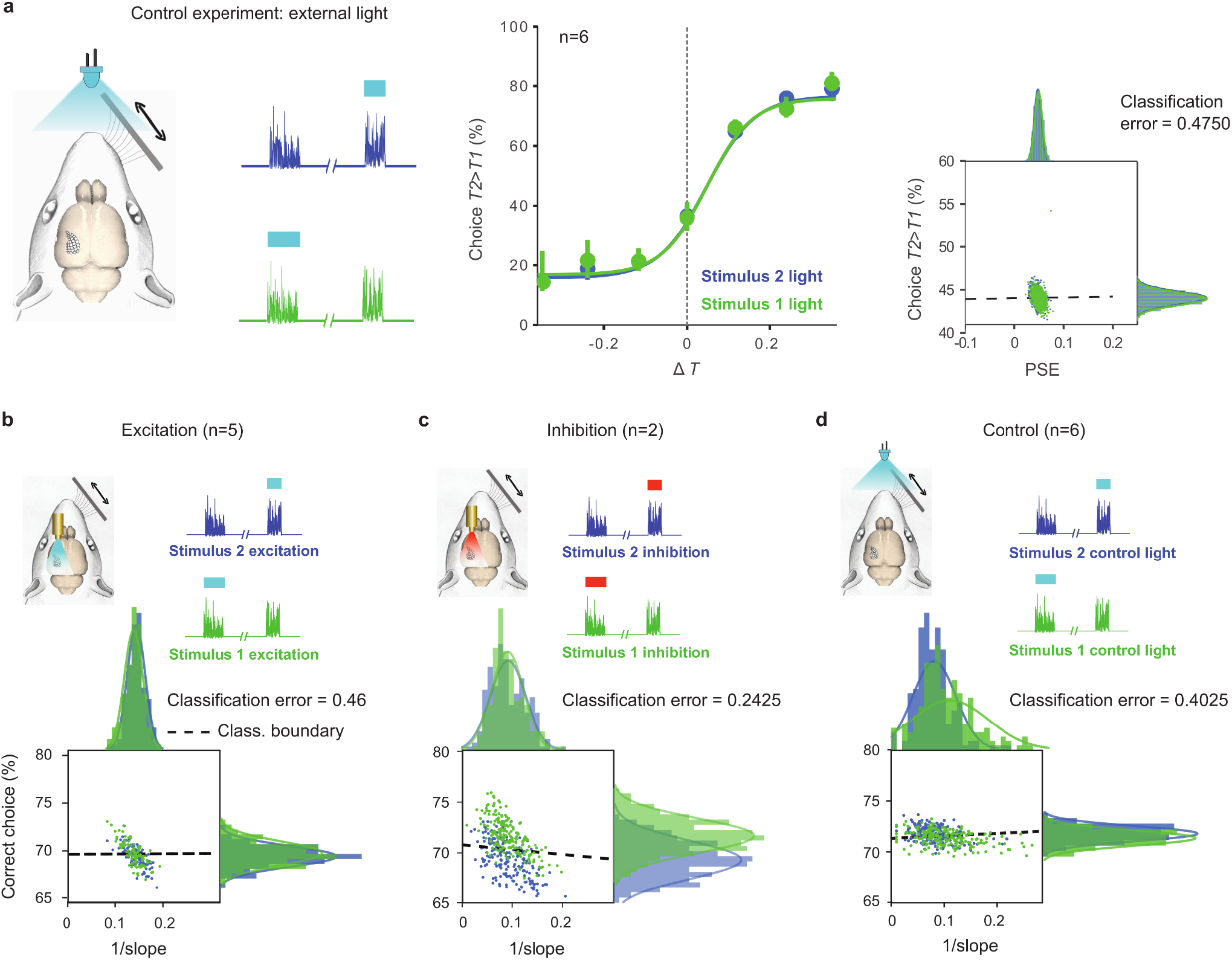
Control experiments. These combine vibrissal stimulation with external light presentation and report statistical tests to probe for a possible effect of optogenetic excitation and inhibition on duration discrimination acuity. **a**, Left: control condition in which an external LED (465 nm) was illuminated above the rat, in correspondence to optogenetic light delivery during vibrissal stimulation (left vibrissae not shown). Middle: light-on during stimulus 2, compared to light-on during stimulus 1 (green) revealed that perceived duration is not biased by visual cues. Right: two signatures of a possible curve shift, percent of trials judged as *T2*>*T1* irrespective of stimulus duration (ordinate) and PSE, the point of subjective equality (abscissa), were measured with bootstrap resampling methods. A support vector machine classifier quantifies data separation by classification error. **b**, Effect of excitation of vS1 neuronal population on sensory acuity. Two signatures of the acuity, percent correct (ordinate), and the psychometric curve’s slope inverse values (abscissa) were measured with bootstrap resampling method. A support vector machine classifier quantifies the separation in the data. Light-on during the second (blue) did not significantly alter acuity as compared to light-on during stimulus 1 (green). **c**, And **d**, same as **b**, but for optogenetic inhibition and control sessions, respectively.

**Extended Data Figure 3:**
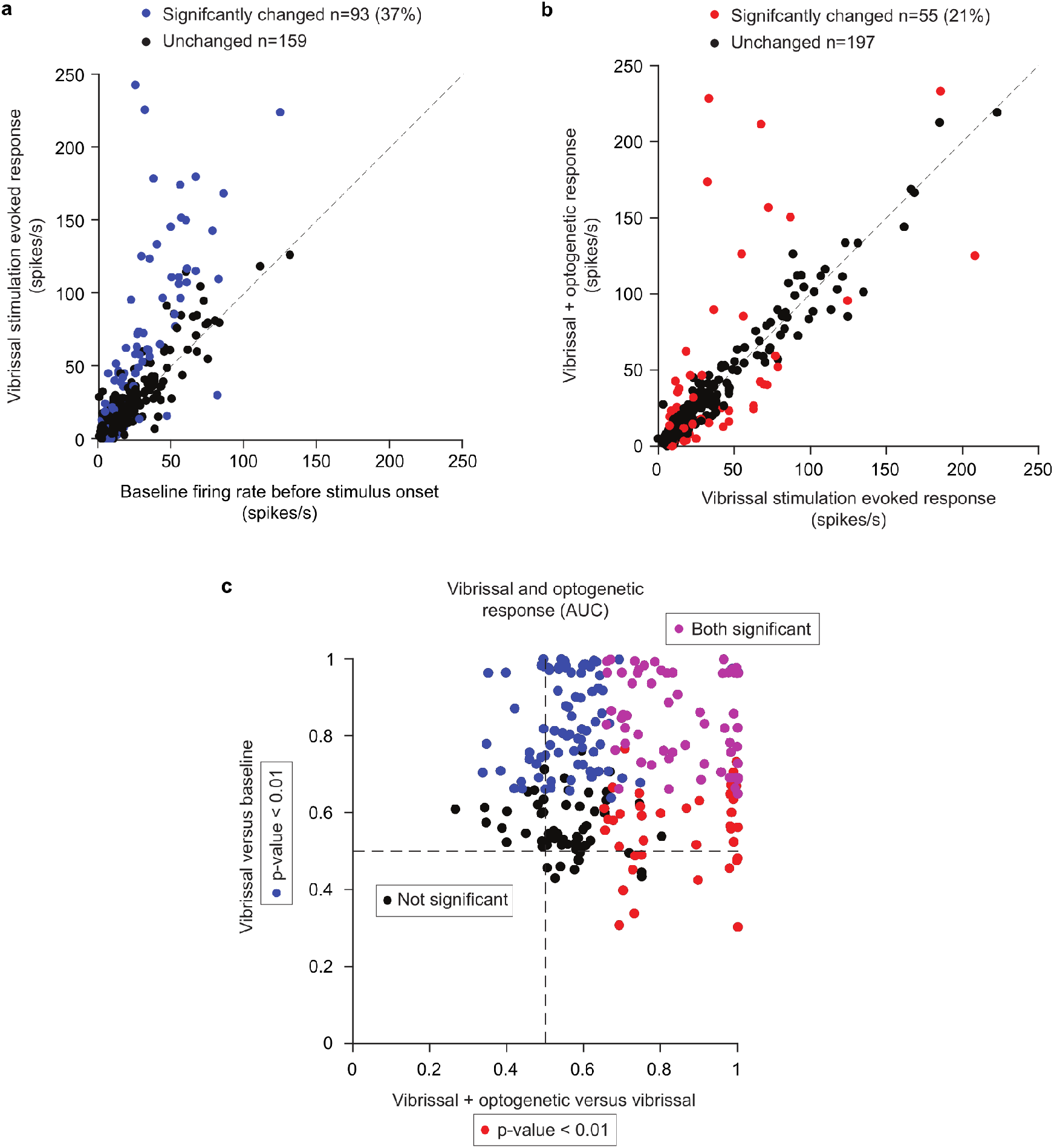
Neuronal responses in vS1 to vibrissal stimulation and optogenetic excitation. **a**, Average firing rate of individual vS1 units before stimulus onset (200 ms time window) versus the first 200 ms during vibrissal stimulation. Blue dots are neurons with significantly (p-value < 0.01, resampling method: permutation test) altered responses by vibrissal stimulation compared to background activity. **b**, The responses of all vS1 units are represented for the first 200 ms of vibrissal stimulation (abscissa), compared to the first 200 ms of vibrissal stimulation accompanied by optogenetic excitation (ordinate). Red dots show neurons with significantly (p-value < 0.01) increased firing rate during optogenetic + vibrissal stimulation as compared to vibrissal stimulation alone. **c**, Area under the ROC curve (AUC) was measured by comparing the response of neurons during the background activity versus vibrissal stimulation (ordinate). Additionally, the response during the vibrissal stimulation is compared to optogenetic + vibrissal stimulation (abscissa). Blue and red dots are neurons with significant response AUC for vibrissal stimulation only versus background activity and vibrissal stimulation versus optogenetic + vibrissal stimulation, respectively. Purple dots represent neurons with significant AUC for both comparisons. Significance was tested by the resampling method (500x, permutation test).

**Extended Data Figure 4:**
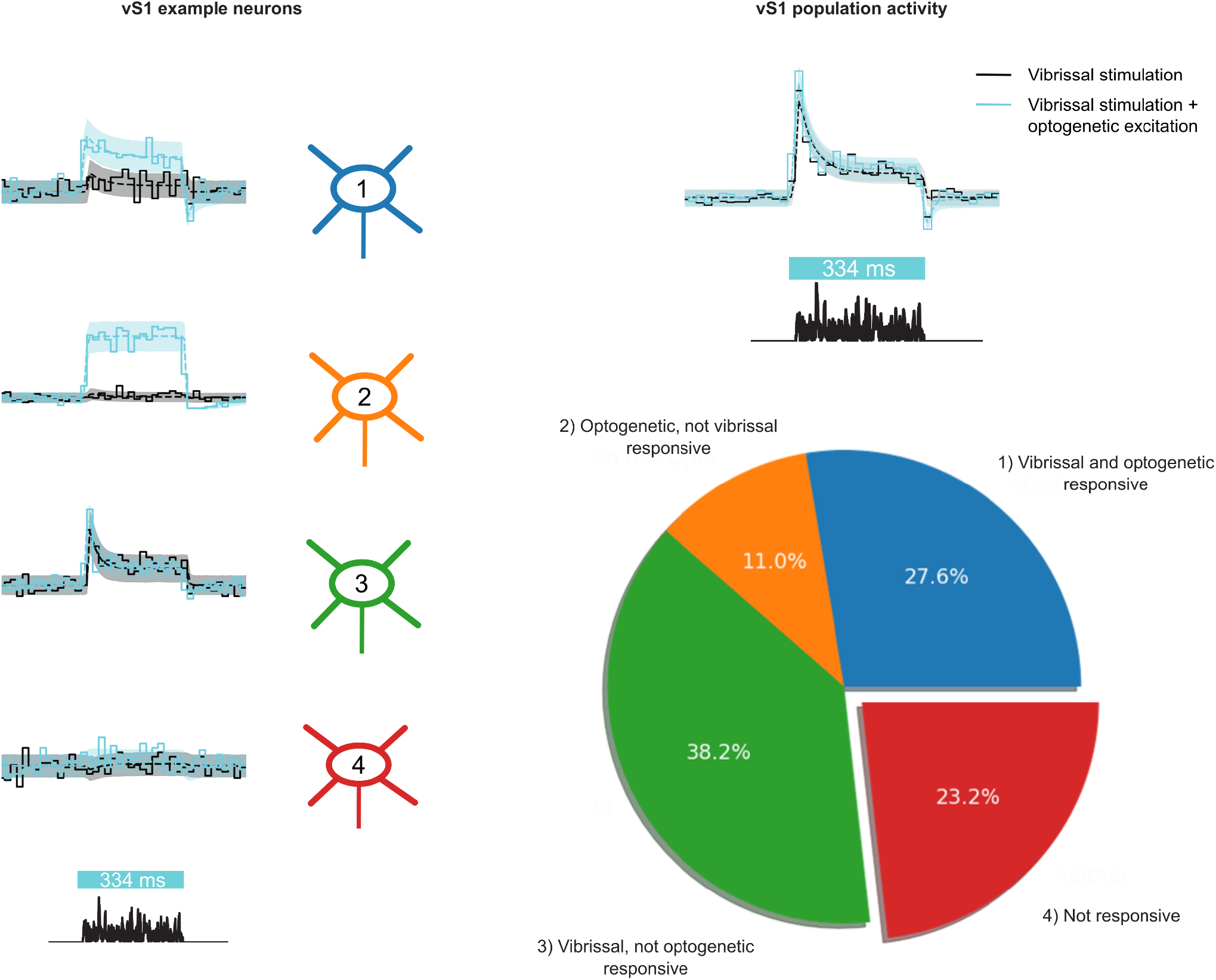
Single vS1 neuron and population responses can be simulated by fitting input current parameters; classification of vS1 neurons. Left: PSTHs of four example neurons, responding to a 334 ms vibrissal stimulation (black, solid lines) and vibrissal stimulation + optogenetic excitation (light blue, solid lines). Neuron-specific response characteristics, based on vibrissal and optogenetic drive, was simulated (dashed lines) by fitting a parameter set (see Eqs. 6-8) that determines the input currents for generating Poisson spike trains. Confidence intervals (shaded area) of the simulated PSTHs covered the variability of the data. Assessing the fitted parameters for significant (p-value < 0.005) deviation from zero for vibrissal and optogenetic driven currents, allowed a neuron classification based on response properties (pie chart).

**Extended Data Figure 5:**
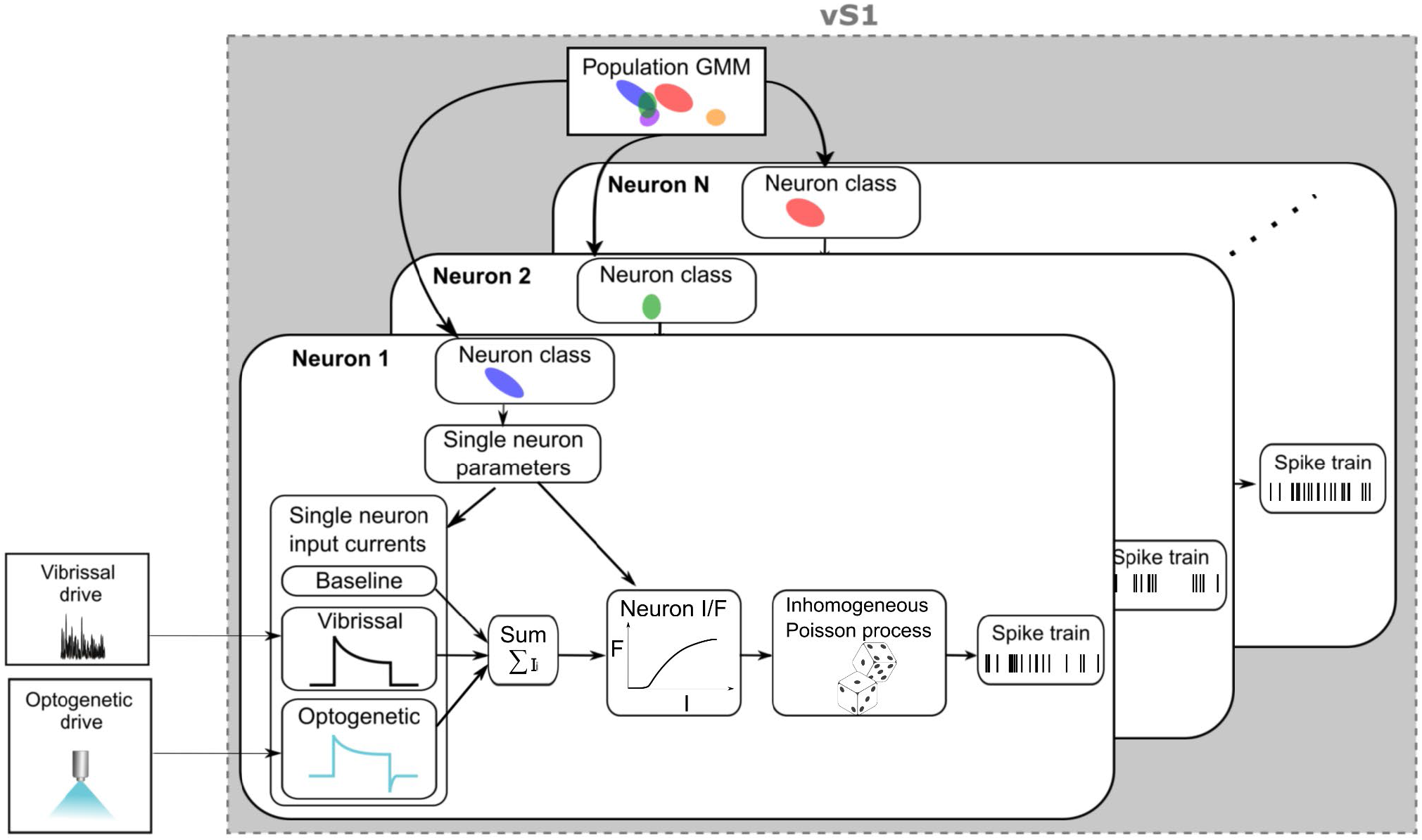
Gaussian mixture model (GMM) of vS1 neuronal activity. Single vS1 spike trains in response to vibrissal stimulation and optogenetic excitation (see leftmost boxes) could be simulated by applying dynamic input currents to a Poisson process through a sigmoidal IF-curve (Eq. 10). The parameter set (see Eqs. 6-8) that determines the input currents and its dynamics, depending on vibrissal and optogenetic drive, was fitted to each recorded neuron individually. The distributions of this parameter set across all fitted neurons did not reveal distinct response-specific clusters and served as representative vS1 population characteristics. The GMM estimated a Gaussian distribution for each of the given parameters from the parameter values, given by fitting each individual neuron. Applying the GMM, representative vS1 model neurons could be resampled (see e.g., *Neuron 1* box), by retrieving a given input current parameter set from the parameters Gaussian distributions. Each model neuron consisted of a distinctive parameter set, representative to the recorded vS1 population that determined the input current, depending on vibrissal and optogenetic drive. Poisson spike trains of resampled model neurons were generated to simulate a vS1 population response (gray box) to a given external input.

**Extended Data Table 1:** Model fitted parameters.

|  |  |  |
| --- | --- | --- |
| $\tau$ | Leaky integration time constant | 990 ms |
| $\mu_b$ | Mean drive excluding vS1 | 1.95 |
| $\sigma_b^2$ | Variance of drive excluding vS1 | 208 |
| $p_L$ | Probability of lapse trial | 0.17 |
| $b_L$ | Probability of choosing $T2 > T1$ in lapse trial | 0.514 |

